# Indel driven rapid evolution of core nuclear pore protein gene promoters

**DOI:** 10.1101/2023.01.04.522740

**Authors:** David W. J. Mcquarrie, Adam M. Read, Frannie H. S. Stephens, Alberto Civetta, Matthias Soller

## Abstract

Nuclear pore proteins (Nups) prominently are among the few genes linked to speciation from hybrid incompatibility in *Drosophila*. It was previously found that neuronal wiring underlying the female post-mating response induced by male-derived sex-peptide requires channel Nup54 functionality. A hot spot for rapid evolution in the promoter of *Nup54* suggests a critical role for regulatory elements at the onset of speciation. Systematic analysis of Nup coding and promoter regions using *Drosophila* phylogenomics reveals that polymorphism differences between closely related *Drosophila* species in Nup coding regions do not generally evolve rapidly. Consistent with findings for *Nup54*, additional channel Nups 58 and 62 promotors are also hotpots for rapid accumulation of insertions/deletions (indels). Examination of Nup upstream regions reveals that core nuclear pore complex gene promoters accumulate indels rapidly. Since changes in promoters can have dominant effects (effects which directly impact gene expression of associated genes), these results indicate an evolutionary mechanism driven by indel accumulation in core Nup promoters. Compensation of such deleterious changes could lead to altered neuronal wiring, rapid fixation of adaptive traits and subsequently the rise of new species. Hence, the nuclear pore complex may act as a nexus for species-specific changes *via* nucleo-cytoplasmic transport regulated gene expression.

## Introduction

Male derived substances transferred during mating change female physiology and behaviour to guarantee reproductive success in many insects ^1–4^. Rapid divergence in male versus female interactive molecules can hamper reproductive success, potentially leading to the establishment of new species for example by compromising fertility or viability among hybrids ^5–13^. Such hybrid incompatibility among closely related *Drosophila* species has been used to identify responsible genes. Among the few genes identified were nuclear pore proteins Nup98 and Nup160 ^14–17^. In addition, a screen for sex-peptide insensitive mutants identified Nup54 functionality to be important for neuronal wiring of circuits involved in regulating post-mating behaviours ^10^. In particular, a deletion in the promoter of the *Nup54* gene has been associated with altered nucleo-cytoplasmic shuttling in eight *pickpocket* (*ppk*) expressing neurons in the central brain leading to wiring defects and a compromised female post-mating response directed by male-derived sex-peptide transferred during mating ^10^. Moreover, this *Nup54* promoter deletion allele maps to a hot spot for rapid evolution in the *Nup54* promotor suggesting sexual conflict driving female escape from male manipulation by sex-peptide under unfavourable conditions ^10^. Furthermore, channel Nups have also been attributed a role in transposon silencing in the germline by facilitating processing of short piRNAs from long pre-curser RNAs of the *flamenco* locus, which functions as a ‘master off switch’ for transposons ^18^, indicating that the disruption to NPC regulation likely has pleiotropic effects.

The nuclear pore protein complex (NPC) provides a physical barrier between nucleus and cytoplasm requiring active transport for cargos above about 40 kDa ^19^. The NPC consists of 30 evolutionary highly conserved proteins in *Drosophila* (Fig. 1), which are grouped into subcomplexes termed outer ring (OR), inner ring (IR), cytoplasmic filaments (CF), nuclear basket (NB) and pore membrane proteins (POMS). The transport channel is made up of three phenylalanine-glycine-rich (FG) repeat domains of Nup54, Nup58 and Nup62. The NPC can contribute to differential expression of other genes as e.g. actively transcribed genes can be in the proximity of the NPC ^20–23^.

**Figure 1:**
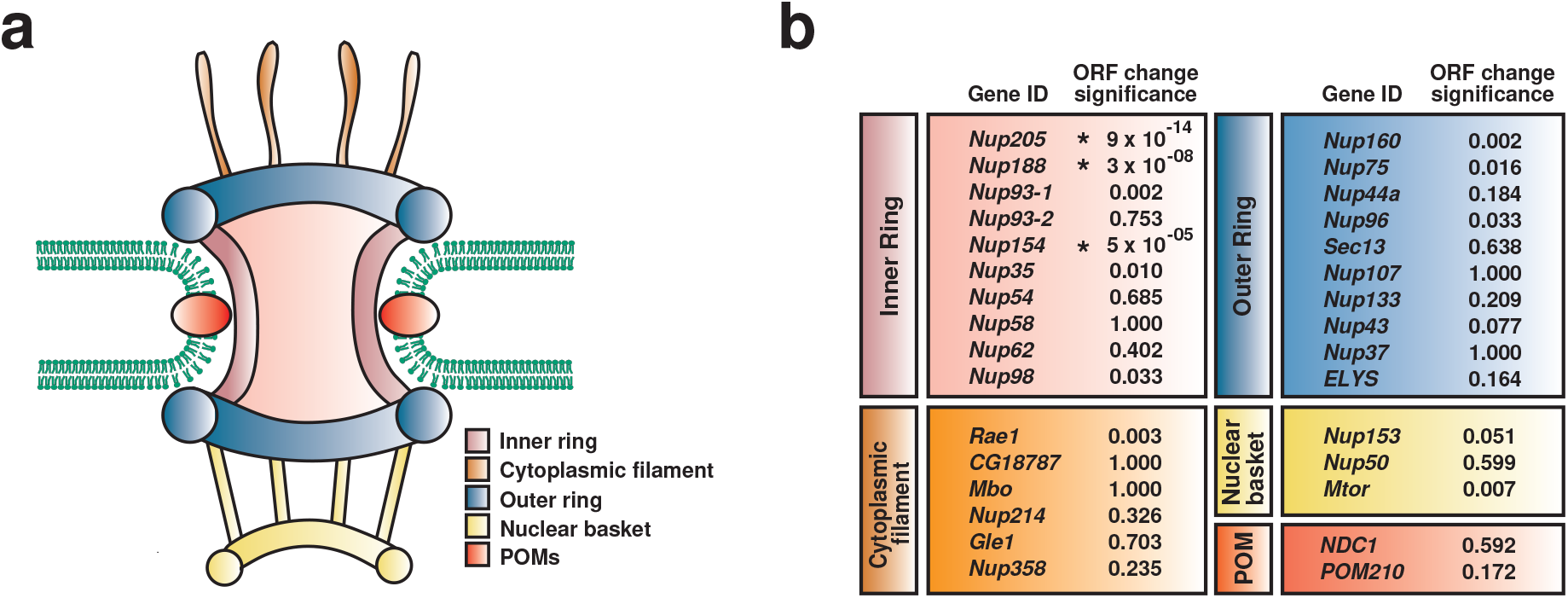
Core nuclear pore protein genes do not generally evolve through adaptation in coding sequences (ORFs). **a**) Graphic depiction of the nuclear pore complex with differentially coloured subcomplexes inner ring (IR, pink), outer ring (OR, blue), cytoplasmic filament (CF, orange), nuclear basket (yellow) and pore membrane proteins (POMS, red) embedded in the nuclear envelope lipid-bilayer (green). **b)** Significance of synonymous to non-synonymous changes in open reading frames (ORFs) indicated by an asterisk and p-values.

Given the rapid evolution of *Nup54* gene expression control and the links of Nups to speciation, we systematically examined, whether the NPC is under increased evolutionary constraint as the molecular mechanisms of how the NPC is involved in speciation remains unexplained. Our systematic analysis of Nup coding and promoter regions using *Drosophila* phylogenomics reveals promoters of Nup core genes as hot spots for insertion/deletion (indel) driven rapid evolution and the nuclear pore complex as a nexus for species-specific changes *via* nucleo-cytoplasmic transport regulated gene expression.

## Results

### Rapid evolution of core nuclear pore protein genes is not generally driven through adaptation in coding sequences

To assess whether the protein coding region (Open Reading Frame, ORF) of Nups were under selection, we performed McDonald-Kreitman tests (MKTs) to compare the number of polymorphisms in the *D. melanogaster* ancestral Congo population, and between *D. melanogaster* and its closest relative *D. simulans*. This analysis showed that most Nup ORFs do not deviate from neutral evolution (Fig. 1b), but that 30% (3 out of 10) of the Nups constituting the IR showed significant evidence of selection.

### Promoter regions of channel Nups 58 and 62 have diverged in closely related species

Since we previously identified the promoter of the channel Nup54 as a hot spot for rapid evolution ^10^, we examined the promoters of the other two channel Nups 58 and 62 among closely related species of *Drosophila* (*D. melanogaster*, *D. simulans*, *D. sechellia*, *D. yakuba* and *D. erecta*) (Fig. 2). This analysis revealed that also the promoters of the other two channel Nups are hot spots for rapid evolution (Fig. 2). Moreover, these changes contain a number of indels (Fig. 2c and d) that can fundamentally impact on transcription factor binding commonly found in the proximal region before the TATA box ^24–28^.

**Figure 2:**
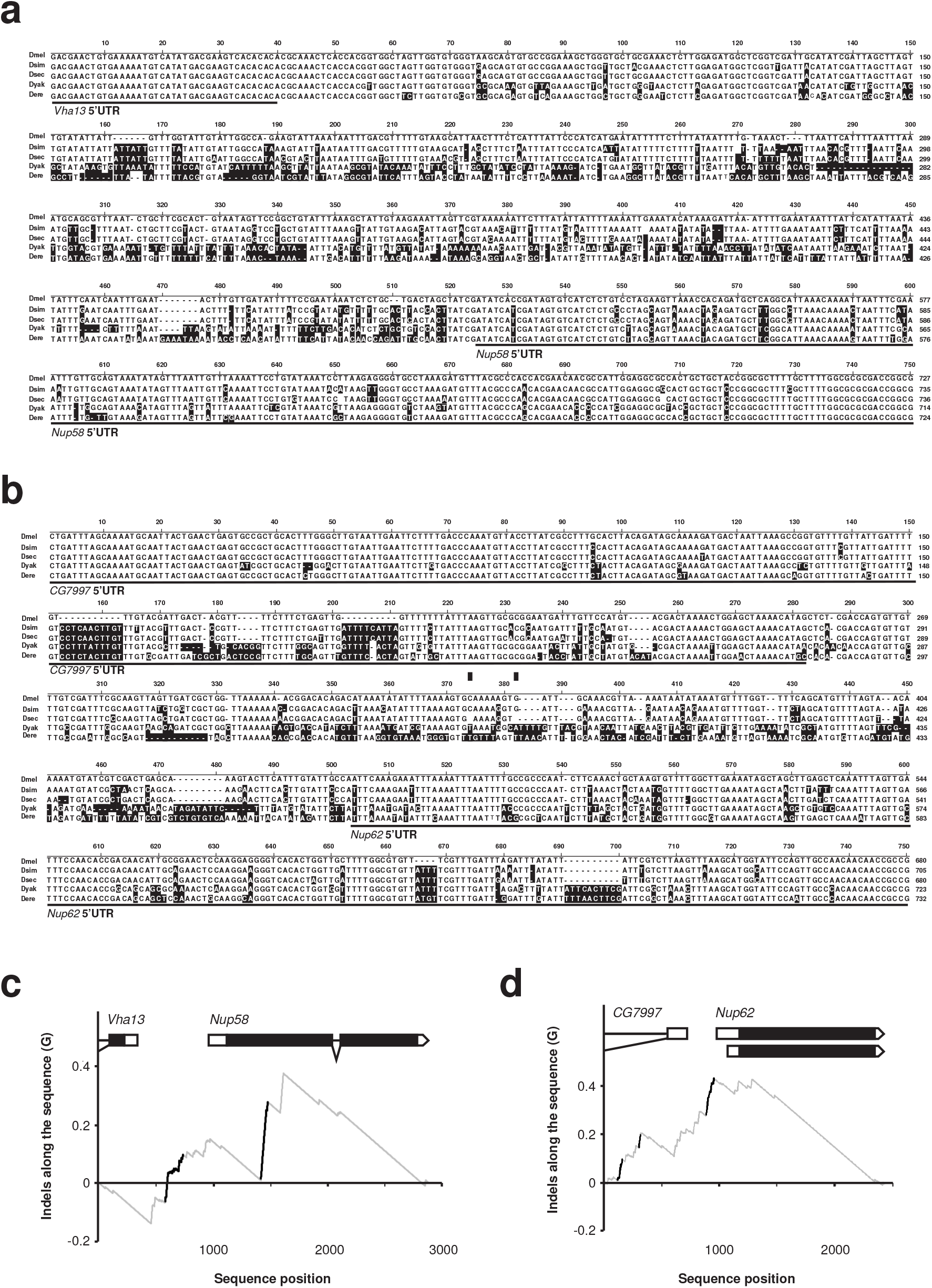
Promoter regions of channel Nups 58 and 62 have diverged in closely related species. **a, b)** Sequence alignment of the Nup58 and Nup62 promoter regions from closely related species. Nucleic acids changes from *D. melanogaster* are indicated in black. Transcribed parts of the *Vha13* and *CG7997* 3’UTR and the *Nup58* and *Nup62* 5’UTR are indicated by a line. **c, d)** Plot of cumulative differences along the sequence (G) between the relative occurrences of indels and their position from the alignment of the gene region around *Nup58* and *Nup62* between *D. melanogaster*, *D. simulans*, *D. sechellia*, *D. yakuba* and *D. erecta*. Positions in the alignment with significant stretches of substitutions are identified by black line(s).

### Inner and outer ring nuclear pore protein gene subcomplexes undergo rapid evolution through indel accumulation in promoters

Next, we examined the promoter region of all remaining Nups in the closely related *Drosophila* species to see whether accumulation of mutations is a general feature of the promoters of this class of genes (Fig. 3, Supplementary Fig. S1). We analysed conservation of in 27 insect species through the PhyloP27way data ^29,30^. To determine whether the NPC evolves rapidly compared to the genome, we used PhyloP scores as a measure of conservation upstream of the TSS (Fig. 3a). We observed a significant decrease in NPC sequence conservation in the promoter region upstream of the predicted TATA box site (−30 to −380) (Fig. 3b and c). To focus more specifically on *Drosophila*, we analysed promoter evolution between species in the *melanogaster* subgroup ^31^. As a control group we used the m^6^A mRNA methylation machinery (consisting of Mettl3, Mettl14, fl(2)d, virilizer, flacc, nito and Hakai and the two readers YTHDC1 and YTHDF), because high evolutionary conservation is required to maintain complex stoichiometry to guarantee functionality, making this group optimal for use as a control group to monitor promoter evolution ^32–35^. In contrast to the genes from the m^6^A mRNA methylation machinery, promoters of ribosomal genes are different in their transcription start site (TSS) properties ^36,37^.

**Figure 3:**
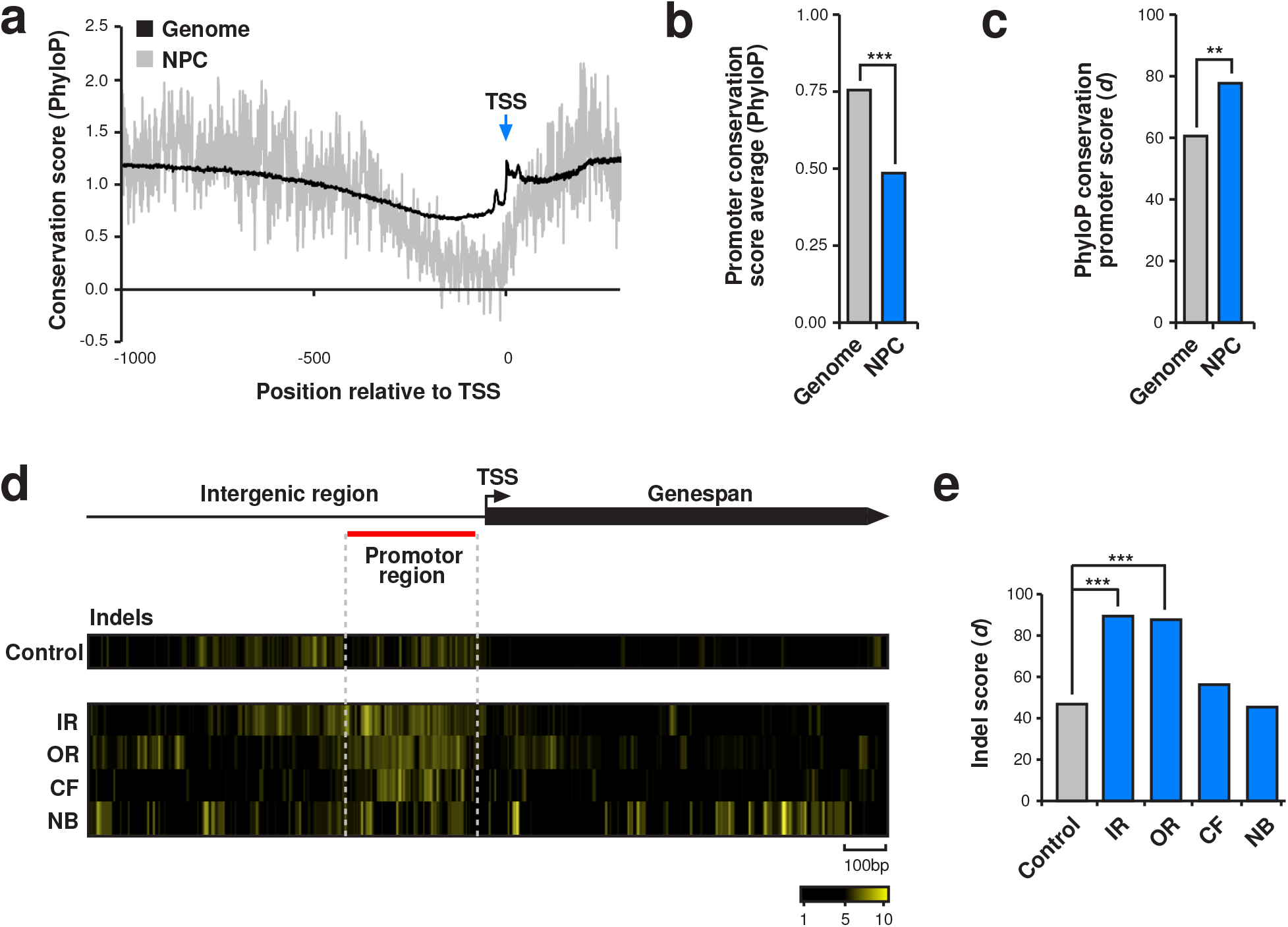
Core nuclear pore protein genes rapidly evolve through indel accumulation in promoters. **a)** Evolutionary conservation of nucleotide positions around the TSS depicted by PhyloP27way conservation score averages for the *D. melanogaster* genome (black) and the NPC (grey). **b)** Evolutionary conservation measured by PhyloP27way conservation score averages for the 350-nucleotide promoter regions compared between the NPC and the genome. Statistically significant differences from unpaired student t-tests are indicated by asterisks (* p≤0.05, ** p≤0.001, *** p≤0.0001). **c)** Evolutionary conservation measured by PhyloP27way conservation promoter *d* scores for the 350-nucleotide for the promoters of the NPC and the genome averages. Statistically significant differences from non-parametric chi-squared tests are indicated by asterisks (* p≤0.05, ** p≤0.001, *** p≤0.0001). **d)** Heatmaps indicating divergence for indels in yellow among closely related *D. melanogaster*, *D. simulans*, *D. sechellia*, *D. yakuba* and *D. erecta* for the control group of genes (m^6^A writer complex and readers) and the inner ring (IR), outer ring (OR), cytoplasmic filament (CF) and nuclear basket (NB) below the gene model with the transcription start site (TSS) indicated by an arrow. The red line indicates the promoter region used for quantification of the substitution rate. **e)** Quantification of the change rate of indels in the TATA box distal region (from −30 to −380). Statistically significant differences from non-parametric chi-squared tests are indicated by asterisks (* p≤0.05, ** p≤0.001, *** p≤0.0001).

As has been observed for *Drosophila* and human promoters ^28,36,38^, the region immediately before the TATA box constitutes transcription factor binding sites (−30 to −380), and showed a slightly increased rate of sequence changes in m^6^A mRNA methylation writer complex and readers consistent with the general trend (Fig. 3d and e, Supplementary Fig. 1). We then analysed the promoters of the different Nup sub-complexes IR, OR, CF and NF. This analysis revealed that the core nuclear complex consisting of the IR and OR genes showed a significantly enhanced rate of indel driven evolution compared to the control group or the CF and NB group of genes (Fig. 3 and Supplementary Fig. S1c-f). We next analysed all sequence differences and performed a more detailed analysis split into deletions, insertions and base changes, relative to *D. melanogaster*. Analysis of all sequence differences reflected our indel analysis, with a significant increase in the number of differences in IR and OR Nups (Supplementary Fig. S1a and b). Our more detailed analysis revealed a significant increase in the number of deletions in IR, OR and to a lesser extend in the cytoplasmic filaments (CF) compared to the control (Supplementary Fig. S1c and d), while no significant changes were detected for insertions (Supplementary Fig. S1e and f). In contrast base changes are increased for IR and reduced for OR and CF (Supplementary Fig. S1g and h).

## Discussion

Here, through comparison of polymorphism rates between closely related Drosophila species we show that rapid evolution of Nups is not generally driven through changes in ORFs. Focussing on upstream and downstream Nup gene regions, we reveal a bias for inclusion of nucleotide changes in the core members of the NPC. Consistent with channel *Nup54* ^10^, further analysis of upstream regions of the other channel Nups 58 and 62 reveals their promotor regions to be hotspots for rapid indel accumulation. Through examination of promoter regions of Nup subcomplex members, we identify the core IR and OR Nups as hot spots for rapid accumulation of indels. Here, the core NPC members likely act as drivers of species-specific variation through dominant indel driven changes in promoters, subsequently modifying regulation of nucleo-cytoplasmic transport regulated gene expression.

Although the megadalton nuclear pore protein complex (NPC) has mainly been associated with providing a physical barrier between nucleus and cytoplasm, recent discovery of pleiotropic functions of channel Nup54 provide new insights how the NPC could drive speciation. Nup54 is required for neuronal wiring underlying the female post-mating response induced by male-derived sex-peptide. Since its promoter is a hot spot for rapid evolution, a critical role for regulatory elements at the onset of speciation is indicated as a result of sexual conflict ^5,9,10,13,39^.

Given the role of channel Nups in piRNA meditated transposon silencing of the *flamenco* locus in the germline and their requirement for sexual differentiation ^10,18^, changes at one trait could cascade into the second leading to rapid phenotypic evolution and speciation. Intriguingly, mutations in *cis*-regulatory sequences of genes are often dominant whereas the majority of coding mutations are recessive ^40–42^ leading to different efficacy of selection ^43–45^.

Maternally inherited piRNAs are essential to transposon silencing and an imbalance can lead to a phenomenon called hybrid dysgenesis, imposing reduced fecundity in the female offspring when the male genome contributes novel transposable elements to be silenced ^46^. Key to silencing are epigenetically inherited piRNAs from the female to prime the “ping-pong” cycle for amplification to increase the silencing capacity by heterochromatinization for preventing transposon mobilisation ^46–49^. Compromised transposon silencing leading to deleterious reductions of fecundity could be escaped by altered NPC function. Such changes are likely to trigger pleiotropic effects including altered neuronal wiring and change of behaviour, leading to accelerated evolution and speciation. In essence, if escape from lower fecundity as a result of hybrid dysgenesis ^46–49^, leading to reduced transposon silencing is coupled to changes in the morphology or behaviour, new species could be established very quickly. In particular, if changes in Nup functionality result in rewiring of neuronal circuits as indicated for Nup54, behavioural preferences could lead to rapid isolation. In fact, mate preference has changed in *D. simulans* through rewiring of sensory neuron projections to fruitless P1 neurons that control courtship ^50^.

## Conclusion

Changes in gene expression have profound effects during species divergence and phenotypic adaptation, and such changes can lead to hybrids’ gene mis-expression and dysfunction ^51–55^. The molecular mechanism how new species arise through differential regulation of gene expression remains uncertain. The newly discovered role of channel Nups in piRNA processing in the germline to maintain transposon silencing provokes the claim that any compensation of the negative impact on fecundity from hybrid dysgenesis would be favoured. However, due to the pleiotropic effect of Nups on neuronal wiring and behaviour, the escape from hybrid incompatibility could produce changes leading to behavioural isolation ^10,18^. Since mutations in promoters can exert a dominant effect ^40–42^, such changes could be rapidly fixed in contrast to recessive changes in coding regions, which would remain hidden in heterozygosity. Our systematic analysis of the evolution of all Nups coding and promoter regions suggests a less studied mode of evolution through changes of sequences upstream of the transcription start site (TSS), particularly in the promoters of core Nups, and supports the possibility that compensation of deleterious changes in the germline can lead to altered neuronal wiring and rapid fixation of adaptive traits.

## Materials and Methods

### Open reading frame analysis

To determine whether the ORFs of nucleoporins are under selection, the PopFly online database (imkt.uab.cat) developed from the Drosophila Genome Nexus project assembling sequence data of around 1100 *D. melanogaster* genomes were used to perform MKTs to analyse polymorphism data from the ancestral Congo population because it is a sub-Saharan population with higher ancestral stability than other populations ^56–58^. Synonymous and non-synonymous polymorphisms within *D. melanogaster* or between *D. melanogaster* and *D. simulans* were obtained and significance determined by Fisher’s Exact Test, with significance defined as p≤0.05 with Bonferroni correction.

### Sequence/data retrieval and alignment

Nucleoporin gene and promoter sequences were retrieved from UCSC genome browser (genome.ucsc.edu) using the Table Browser tool ^29,30^. Alignments were carried out with clustalW within the MEGA11 program ^59^. PhyloP27way data were sourced from UCSC genome browser through the Table Browser tool ^29,30^. Data points were collected for a region of 1000 nucleotides upstream and 300 nucleotides downstream.

### Accumulation of substitutions along extended gene regions

To test for nonrandom accumulation of substitutions along the Nup extended gene regions we determined significant deviations from a uniform distribution of substitutions using an empirical cumulative distribution function. The function (G) detects monotonic increases in substitutions (n) measured as the difference between the relative occurrence of a nucleotide change and its relative position in the alignment ^60^. Whether differences between the values of the G function (ΔG) between substitutional events deviates from a random accumulation of changes is tested using Monte Carlo simulations to produce 100,000 samples of n events by sampling sites without replacement along the alignment ^60^.

### Comparison of substitution rates

To analyse the promoters of Nups and the control group of the m^6^A writer complex and readers a region of 1000 bp upstream and downstream of the transcriptional start site (TSS) was used with TSS as an anchor point. Regions of differential alignment between the *Drosophila* species were averaged across the NPC subcomplexes, where alignments were translated into events ^61^. Per ^61^, 0 signified no sequence difference between all analysed species and 1 signified comparative sequence divergence for ≥1 species. A sliding window of five nucleotides upstream and downstream of each position was summed and processed to define sliding event (*Se*) scores which were used to generate heatmaps. To calculate sequence change accumulation (percentage of events greater than the average control promoter sliding window score (*d*)), a 350-nucleotide region upstream of the approximated TATA box region was taken. The total number of *Se* scores greater than the control *Se* (*Se^C^*) was divided by the total number of events in the region (*N*).

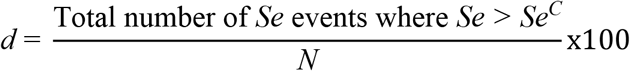

Significance was defined by non-parametric chi-squared tests vs the control score. P values ≤0.05 were deemed statistically significant.

To compare the conservation rate of the NPC compared to the *D. melanogaster* genome, PhyloP27way scores for a region of 1000 nucleotides upstream and 300 nucleotides downstream of the TSS for all *Drosophila* genes was performed. Genes without data points for the full 1300 nucleotide region were omitted. Genome-wide and NPC mean PhyloP scores were calculated using R version 4.4.2 ^62^. The average 350-nucleotide region upstream of the approximated TATA box region was compared between the two and significance was calculated using an unpaired student t-test. P values ≤0.05 were deemed statistically significant. For the same region of the NPC and genome averages *d* was calculated and significance determined as described.

## Data availability

All data generated or analysed during this study are included in Supplementary Dataset S1.

## Acknowledgements

This work was supported by the Medical Research Council Integrated Midlands Partnership for Biomedical Training Program by D.W.J.M., the Biotechnology and Biological Science Research Council to M.S. and the Natural Sciences and Engineering Research Council of Canada to A.C. We thank R. Arnold for help with bioinformatics.

## Authors contributions

M.S. conceived and directed the project. All authors analysed data. M.S., D.W.J.M. and A.C. wrote the manuscript. All authors read and approved the final manuscript.

## Additional information

Supplementary information includes Fig. S1 and Supplementary Dataset 1.

## Figure legends

**Supplementary Figure S1:**
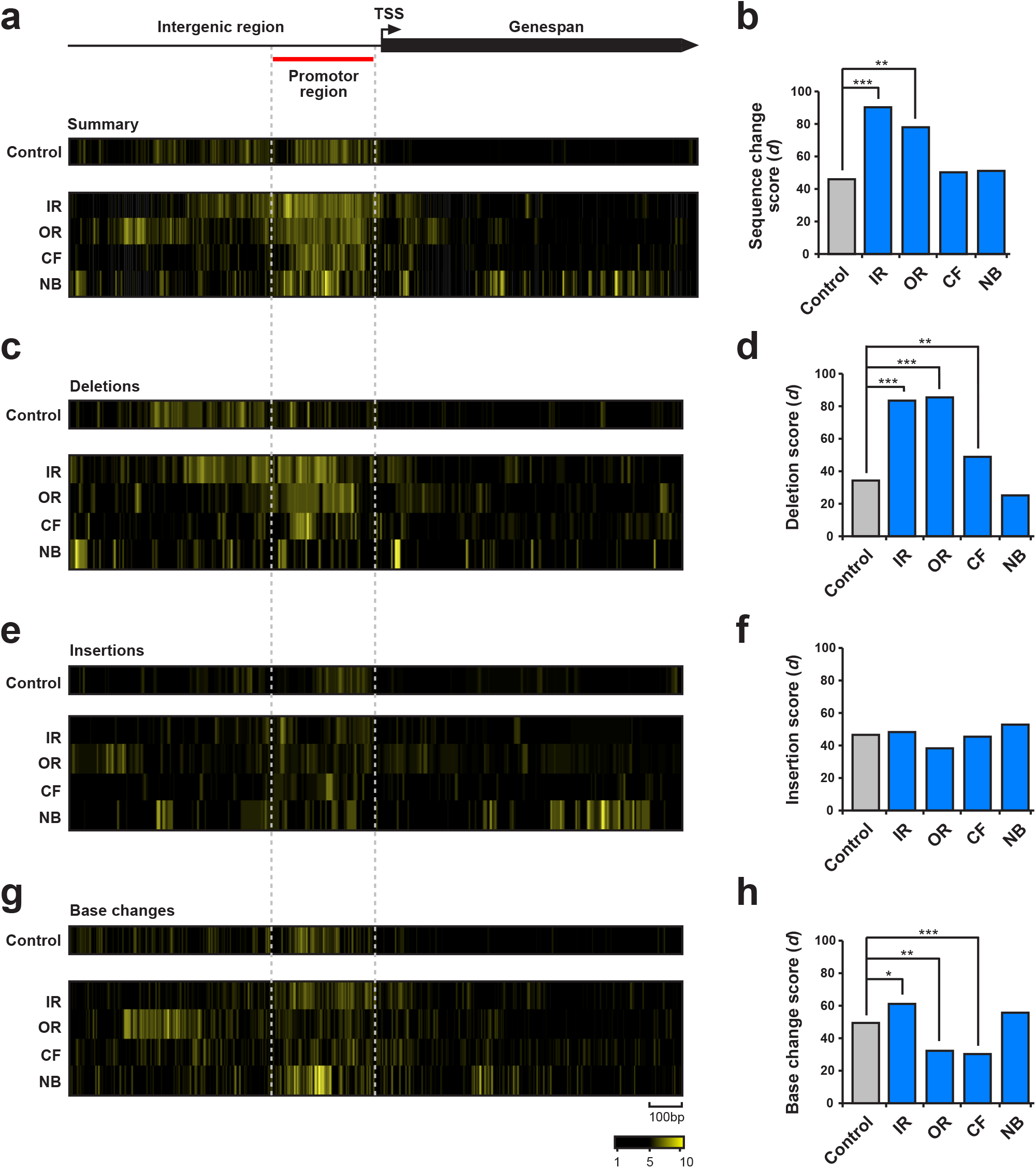
Inner and outer ring nuclear pore protein gene subcomplexes undergo rapid evolution through indel accumulation in promoters. **a)** Gene model depicting the intergenic region followed by the transcription start site (TSS) and the genespan with the 350-nucleotide region analysed for promotor activity upstream of the TATA box location. **a-h)** Heatmaps indicating divergence in yellow among closely related Drosophila species for the control group of genes (m6A writer complex and readers) and the inner ring (IR), outer ring (OR), cytoplasmic filament (CF) and nuclear basket (NB). Calculated sequence difference index, deletion index, insertion index and base change index events are reported as bar charts for the NPC subgroups compared to the control group. Index events were defined by values above the control intergenic region average. Statistically significant differences between the NPC subgroups and the control group were calculated from chi-squared tests and indicated by asterisks (* p≤0.05, ** p≤0.001, *** p≤0.0001).

